# Improving Metagenomic Assemblies Through Data Partitioning: a GC content approach

**DOI:** 10.1101/261784

**Authors:** Fábio Miranda, Cassio Batista, Artur Silva, Jefferson Morais, Nelson Neto, Rommel Ramos

## Abstract

Assembling metagenomic data sequenced by NGS platforms poses significant computational challenges, especially due to large volumes of data, sequencing errors, and variations in size, complexity, diversity and abundance of organisms present in a given metagenome. To overcome these problems, this work proposes an open-source, bioinformatic tool called GCSplit, which partitions metagenomic sequences into subsets using a computationally inexpensive metric: the GC content. Experiments performed on real data show that preprocessing short reads with GCSplit prior to assembly reduces memory consumption and generates higher quality results, such as an increase in the N50 metric and the reduction in both the L50 value and the total number of contigs produced in the assembly. GCSplit is available at https://github.com/mirand863/gcsplit.

## 1 Introduction

Metagenomics consists in determining the collective DNA of microorganisms that coexist as communities in a variety of environments, such as soil, sea and even the human body [1–3]. In a sense, the field of metagenomics transcends the traditional study of genes and genomes, because it allows scientists to investigate all the organisms present in a certain community, thus allowing the possibility to infer the consequences of the presence or absence of certain microbes. For example, sequencing the gastrointestinal microbiota enables the understanding of the role played by microbial organisms in the human health [4].

Nevertheless, second generation sequencing technologies – which belong to the Next Generation Sequencing (NGS), and are still the most widespread technology on the market – are unable to completely sequence the individual genome of each organism that comprises a metagenome. Instead, NGS platforms can sequence only small fragments of DNA from random positions, and the fragments of the different organisms are blended [5]. Hence, one of the fundamental tasks in metagenome analysis is to overlap the short reads in order to obtain longer sequences, denominated *contigs*, with the purpose of reconstructing each individual genome of a metagenome or represent the gene repertoire of a community [6]. This task is referred to as the metagenome assembly problem.

Roughly speaking, metagenome assembly can be done with or without the guidance of a reference genome. The reference assembly can be performed by aligning reads to the genomes of cultivated microbes [7]. However, this method is rather limited because the microbial diversity of most environments extends far beyond what is covered by the reference databases. Consequently, it is necessary to perform *de novo* assembly when reconstructing a metagenome that contains many unknown microorganisms.

Although it seems simple at first glance, the metagenome assembly problem is actually quite complex. Among the several challenges this task arises, there are sequencing errors specific to each platform and the processing of the large volume of data produced by NGS platforms [8]. Moreover, the problem is further complicated by variations on the size of the genomes and also by the complexity, diversity and abundance of each organism present in a microbial community [9]. For these reasons, the metagenome assembly becomes a challenging problem.

To solve all these challenges, either *de novo* assembly can be performed directly by a metagenome assembler, or the short reads can be clustered in advance in order to individually assembly each organism present in the metagenome [10]. The latter approach has the advantage of reducing the computational complexity during the metagenome assembly, because the assembler will process smaller subsets of short reads and, furthermore, it is possible to run the individual assembly of each genome in parallel, since those tasks are independent from each other. The reduction of computational complexity can also be achieved through the previous digital normalization or data partitioning prior to assembly, which reduces the dataset by removing redundant sequences, and divides it into groups of similar reads, respectively [11].

The main focus of this study is the application of the data partitioning method towards the reduction of computational complexity and the improvement of metagenome assembly. The developed computational approach, denominated GCSplit, uses the nucleotide composition of the reads, i.e., the amount of bases A, G, C and T present on DNA sequences. This decision was based on the fact that different organisms or genes that compose metagenomes have distinct GC content and different GC contents will present coverage variation, a metric used by assemblers to reconstruct the genomes, which in turn affects the k-mer selected to perform the sequence assembly based on NGS reads.

The rest of this paper is structured as follows. Related works on digital normalization and data partitioning are discussed in Section 2. Section 3 then presents the proposed algorithm. In Section 4, the impact of the new approach on the performance of the metagenomic assembler metaSPAdes [12] is evaluated through experiments on real data. Finally, Section 5 presents the conclusions and plans for future works.

## 2 Related Work

In the literature there are several studies that attempt to reduce the computational complexity and improve metagenomic assemblies through data preprocessing techniques. The main approaches used are either digital normalization or data partitioning, the latter being the main focus of this article. In this context, the goal of this section is to carry out a bibliographical review of tools that use such methodologies.

Diginorm [13] is a tool that uses the CountMin Sketch data structure to count k-mers, with the purpose of obtaining an estimate of the sequencing coverage; and reducing coverage variation by discarding redundant data. Due to the data structure, this technique keeps a constant memory usage and a linear runtime complexity for the *de novo* assembly in relation to the amount of input data.

Trinity’s *in silico* normalization (TIS) [14], which belongs to the Trinity assembler algorithm package, presents an implementation that computes the median k-mer coverage for all reads of a given dataset. If the median coverage is lower than the desired value, all reads are kept. Otherwise, the reads may be kept with a probability that is equal to the ratio of the desired coverage by the median coverage.

NeatFreq [15] clusters and selects short reads based on the median k-mer frequency. However, the main innovation in the work is the inclusion of methods for the use of paired reads alongside with preferential selection of regions with extremely low coverage. The results achieved indicate that the coverage reduction obtained by NeatFreq increased the processing speed and reduced the memory usage during the *de novo* assembly of bacterial genomes.

ORNA [16] presents a novel and interesting approach that normalizes short reads to the minimum necessary amount in order to preserve important k-mers that connect different regions of the assembly graph. The authors treat data normalization as a set multi-cover problem, and they also have proposed a heuristic algorithm. Their results show that a better normalization was achieved with ORNA, when compared with similar tools. Moreover, the size of the datasets was drastically reduced without a significant loss in the quality of the assemblies.

Pell *et al*. [17] presented a novel data partitioning methodology, in which the main data structure – a probabilistic model called bloom filter – was used to obtain a compact representation for graphs. The authors’ implementation can represent each k-mer using only 4 bits, which was the major factor for achieving a forty-fold memory economy while assembling a soil metagenome.

MetaPrep [18] contains efficient implementations for k-mer counting, parallel sorting, and graph connectivity and partitioning. The developed solution was evaluated in a soil metagenome dataset composed of 223 Gigabases (Gb) distributed in 1.13 billion short reads. As a result of the experiment, MetaPrep took only 14 minutes to process this dataset using just 16 nodes of the NERSC Edison supercomputer. The authors also assessed how MetaPrep can improve the performance of the metagenomic assembler MEGAHIT.

Cleary *et al.* [19] proposed an approach that separates the DNA sequences into partitions considering the biological factor, thus allowing the individual assembly of each genome present in a metagenome. The proposed methodology assumes that the abundance of the genomes present in a sample reflects on their k-mer abundance. The results achieved allowed the partial or almost complete assembly of bacteria whose relative abundance varies to a minimum of 0.00001%.

During the literature review, it was not found any software that performs data partitioning using the information present in the nucleotide composition of short reads sequenced by NGS platforms. Hence, in this work we propose GCSplit, a tool that uses the GC content of the DNA sequences in combination with statistical metrics to partition the dataset. This new approach is promising because it is computationally inexpensive and uses information present in the reads that, as far as we know, has not been used in any other work for data partitioning. Further details about this new algorithm will follow.

## 3 The Proposed Algorithm

GCSplit was implemented in C++ in order to facilitate the communication with the library that performs the parallelization of the critical sections of the algorithm. The software KmerStream [20] and metaSPAdes [12], which are automatically executed to estimate the best values of k-mer and to assembly the metagenome, respectively, are dependencies of the proposed algorithm. The object-oriented programming paradigm was used in order to simplify an eventual addition of new assemblers and k-mer estimation programs in the future, since one would only need to implement new classes to interact with the desired programs. Figure 1 summarizes the main steps of the developed algorithm.

**Fig. 1.**
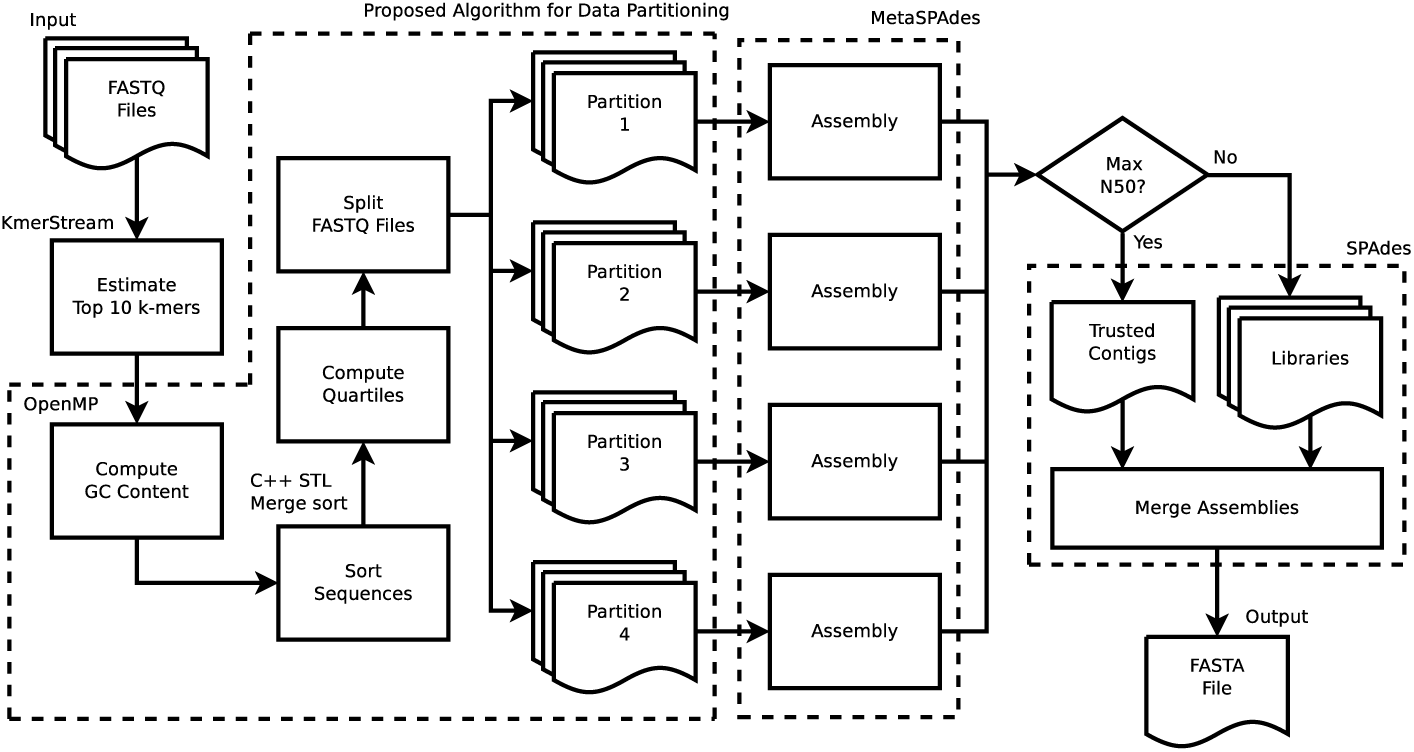
GCSplit algorithm overview.

As input, GCSplit takes two paired FASTQ files, both containing the metagenome’s sequences. The files are automatically passed to KmerStream, which estimates the best 10 k-mer values for the assembly. Next, an algorithm was developed within GCSplit in order to partition the datasets, as shown in the pseudo-code of Algorithm 1. More specifically, the algorithm computes the GC content from all reads, making use of OpenMP library to do so in parallel. C++ STL library, on the other hand, has a parallel implementation of the merge sort algorithm, which is used to sort the sequences according to their GC values in ascending order. The proposed algorithm then calculates the first, second (median) and third quartiles for the set of sorted reads; and divides the dataset into four subsets based on those statistical metrics.

**Algorithm 1:**
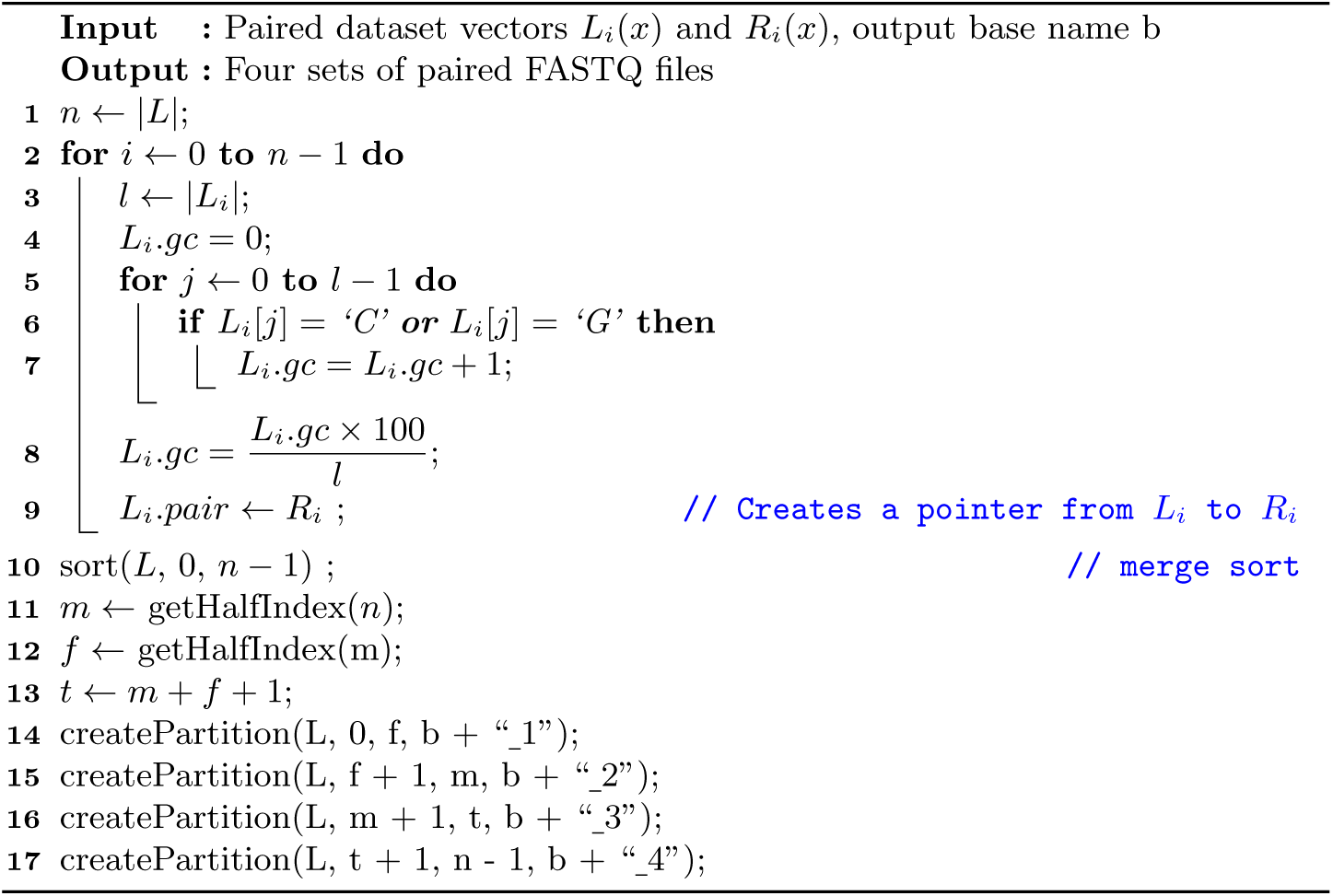
Pseudo-code to partition the dataset

The number of partitions has been set to four because the coverage may become low in an excessively divided dataset and, consequently, the assembly may become fragmented as well. It is important to notice that this partitioning method keeps the reads paired, which is crucial in obtaining assemblies with the highest quality possible.

The calculation of quartiles is performed by the function getHalfIndex, which contains only a conditional test and a simple mathematical operation, thus its complexity can be said to be asymptotically constant, i.e., 𝒪(1). On the other hand, the division of the data into subsets is performed by a procedure called cre-atePartition, whose implementation has four attributions and a single loop.

Therefore, the asymptotic analysis of the second task results in a complexity of 𝒪(*e* – *b*), where “*b*” is the beginning and “*e*” the end of the partition.

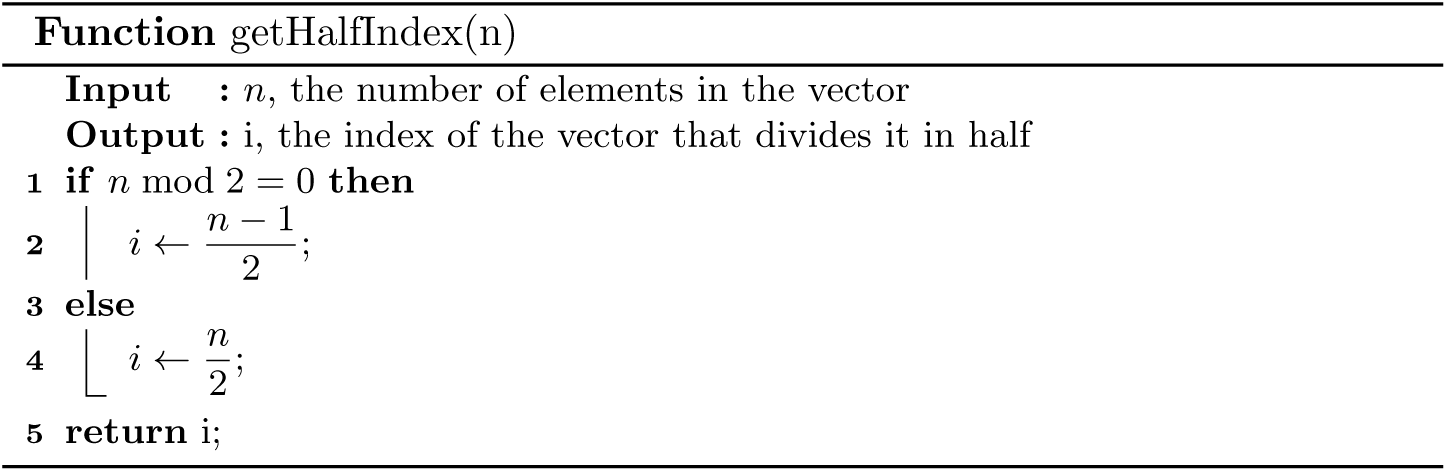

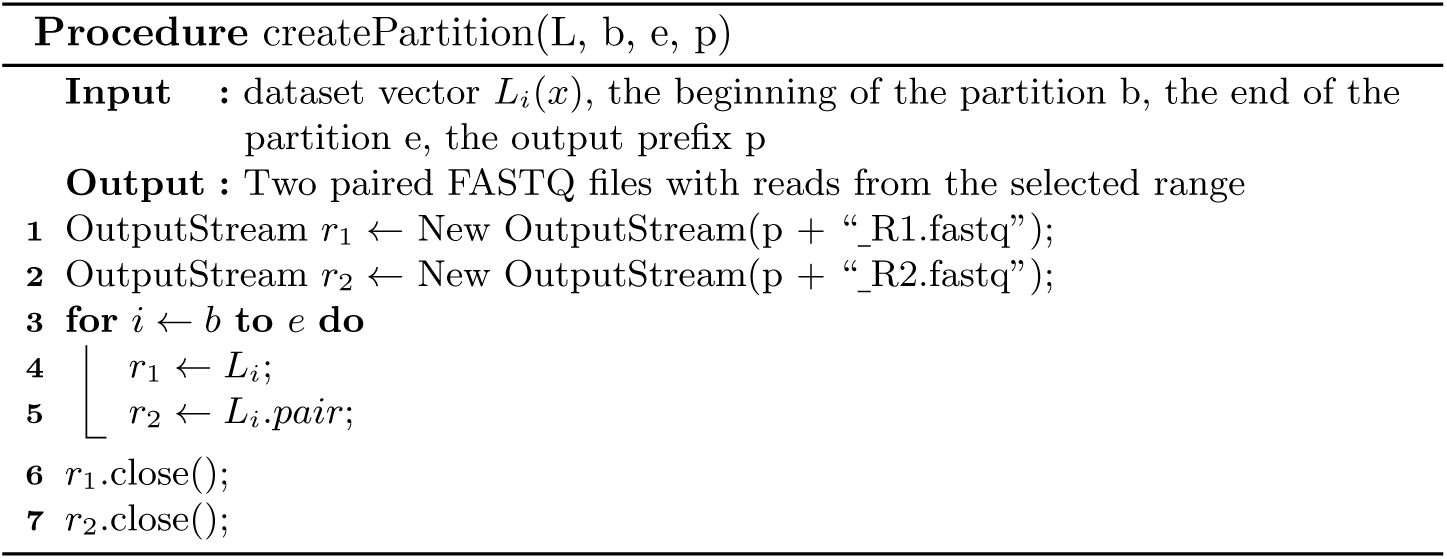

Regarding Algorithm 1, the computation of the number of reads |*L*| takes 𝒪(*n*); sorting with merge sort takes *Θ*(*n* log *n*); calculating the quartiles and dividing the dataset into partitions takes 𝒪(1) + 𝒪(*b – e*); and computing the GC content of “*n*” reads with length “*l*” takes 𝒪(*nl*) due to the nested loop ranging from line 2 to line 9. Therefore, we can conclude that the the proposed partitioning algorithm has an asymptotic complexity of 𝒪(*nl*), since in the worst case *n* x *l* > *n* log *n* > *b –* e.

The created partitions are individually assembled by metaSPAdes. Then, another assembly is performed with SPAdes [21] to concatenate the four previous assemblies, where the assembly result that gets the highest N50 is used as input in the trusted contigs parameter and the remaining are used as single libraries. SPAdes was used in this final step because metaSPAdes does not accept multiple libraries as input yet. Ultimately, the final output is a FASTA file that contains a high quality assembly.

## 4 Evaluation

### 4.1 Datasets and Platform

In order to evaluate GCSplit, analysis were conducted in three real metagenomic datasets, whose samples were collected from the following environments: from a moose rumen, from hot springs at headwaters and a sewage treatment plant. The datasets used in the experiments are listed in Table 1.

**Table 1.** Dataset description

| ID | Dataset | Read Count | Size | SRA Accession |
| --- | --- | --- | --- | --- |
| | | R ($\times 10^6$) | M (Gbp) | Number |
| MR | Moose Rumen | 25.6 | 5.2 | ERR1278073 |
| HS | Hot Springs | 7.4 | 3.7 | ERR975001 |
| STP | Sewage Treatment Plant | 21.3 | 6.4 | SRR2107212,<br>SRR2107213 |

The moose rumen sample was collected in V¨axsj¨o, Sweden. This sample was sequenced with the Illumina HiSeq 2500 platform and contains 25,680, 424 paired short reads with 101 base pairs (bp). These short reads can be obtained from the Sequence Read Archive (SRA) database through the accession number ERR1278073.

The hot springs samples were collected in the years 2014 and 2015 at the headwaters of Little Hot Creek, located in the Long Valley Caldera, near Mammoth Lake in California, United States [22]. Sequencing was conducted using one of the following Illumina platforms: MiSeq PE250, MiSeq PE300 or HiSeq Rapid PE250. The insert size used has approximately 400 bp and MiSeq runs were prepared using the the Agilent SureSelect kit, while HiSeq PE250 samples were prepared using the Nextera XT library preparation kit. These data, which contain 7,422, 611 short reads of 250 base pairs can be obtained via SRA accession number ERR975001.

The sludge sample was collected in a municipal sewage treatment plant located in Argentina [23]. This sample was split in two technical replicates and sequenced with the platform Illumina HiSeq 1500 in 150 bp short reads. This dataset can be downloaded through the SRA accession numbers SRR2107212 and SRR2107213, which contain 8,899, 734 and 12, 406, 582 short reads, respectively.

The assemblies were performed in a cluster that runs the GNU/Linux x64 operating system (openSUSE), and contains 4 nodes with 64 cores and 512 Gigabytes (GB) of Random Access Memory (RAM).

### 4.2 Results

Table 2 shows the assembly quality results produced by metaSPAdes with and without preprocessing for the metagenomes collected from the Moose Rumen (MR), the Hot springs (HS) and the Sewage Treatment Plant (STP) datasets. Assembly statistics were computed with the tool MetaQUAST [24], while peak memory usage was extracted from the assemblers’ log. The best results are highlighted in bold.

**Table 2.** Assembly quality comparison

| Dataset | Preproc.? | MetaSPAdes assembly statistics |  |  |  |  |  |
| --- | --- | --- | --- | --- | --- | --- | --- |
|  |  | #Contigs | Largest Contig | Total Length (Mbp) | N50 | L50 | Memory Peak (GB) |
| MR | Yes | **6,656** | 42,585 | 12.9 | **2,875** | **959** | **19** |
|  | No | 175,829 | **408,484** | **276.1** | 2,003 | 23,838 | 34 |
| HS | Yes | **989** | 101,712 | 4.8 | **12,589** | **86** | **20** |
|  | No | 26,656 | **109,705** | **28.6** | 1,108 | 4,795 | 45 |
| STP | Yes | **11,224** | 114,168 | 28.4 | **5,059** | **1,133** | **33** |
|  | No | 385,566 | **340,186** | **463.8** | 1,312 | 71,126 | 76 |

For the MR dataset, the amount of contigs was drastically reduced from 175, 829 to 6,656 with the previous preprocessing by GCSplit. The N50 value increased from 2, 003 bp in the assembly without preprocessing to 2, 875 bp after preprocessing, while the L50 value decreased from 23, 838 to 959 contigs with the aid of GCSplit. This implies that to reach the N50 value of 2, 875, we need to sum the length of only 959 contigs, whereas without GCSplit 23,838 contigs were necessary to reach approximately the same value, which means that the assembly with metaSPAdes alone contained smaller sequences overall. There was also a reduction of 15 GB in memory consumption during the assembly with GCSplit.

Nevertheless, the largest contig produced in the MR assembly after preprocessing the dataset with GCSplit decreased from 408, 484 bp to 42, 585 bp. Similarly, the total length of the assembly generated after applying GCSplit to the data dropped from 276.1 Mbp to 12.9 Mbp. The merging strategy adopted may be one of the reasons for these reductions, but other possibilities are under investigation.

The assembly of the HS sample also showed improvements with the usage of GCSplit. Memory peak dropped from 45 GB to 20 GB on partitioned data. Furthermore, the amount of contigs decreased from 26, 656 in the assembly with metaSPAdes alone to 989 contigs after preprocessing with GCSplit, representing a 96% reduction. On the other hand, the N50 value increased significantly, yielding a 1036% growth after partitioning, which is excellent for gene prediction. Moreover, the L50 value reduced about 98% with the GC content partitioning.

In the HS dataset, the size of the largest contig produced in the assembly after preprocessing was closer to the assembly with metaSPAdes alone, containing 101, 712 bp and 109, 705 bp, respectively. The total length of the assembly also experienced a less dramatical decrease, going from 28.6 Mbp in the traditional assembly to 4.8 Mbp after using GCSplit.

For the STP sample, there was a 56% economy in memory usage with prior data partitioning. Additionally, the amount of contigs drastically reduced from 385, 566 to 11, 224 contigs when the assembly was performed after preprocessing the sequences with GCSplit. The N50 value raised from 1,312 bp in the assembly with metaSPAdes alone to 5, 059 bp after partitioning, while the L50 value decreased from 71, 126 to 1, 133 contigs with the aid of GCSplit.

However, the largest contig produced in the STP assembly after preprocessing the dataset with GCSplit decreased from 340, 186 bp to 114, 168 bp. Likewise, the total length of the assembly generated after applying GCSplit to the data declined from 463.8 Mbp to only 28.4 Mbp. This result shows that despite the amount of data has decreased, the N50 value improved and this can favor gene prediction in later analysis.

## 5 Conclusion

In this work, we developed a new bioinformatic tool called GCSplit, which partitions metagenomic data into subsets using a computationally inexpensive metric: the GC content of the sequences. GCSplit has been implemented in C++ as an open-source program, which is available at https://github.com/mirand863/gcsplit. It requires GCC version 4.4.7 or higher, the library OpenMP and the software KmerStream and metaSPAdes.

Empirical results showed that applying GCSplit to the data before assembling reduces memory consumption and generates higher quality results, such as increase in the N50 metric and reduction in the L50 value and in the total number of contigs generated in the assembly. Although assemblies with GCSplit produced less data, it is important to notice that the next analysis performed after assembly is gene prediction, where larger sequences are more likely to have genes predicted, as opposed to fragmented assemblies such as those carried out without GCSplit, which contain smaller N50 and larger amounts of bp.

As future work, we will explore alternative merging strategies to improve the size of the largest contigs generated in the assembly. Additionally, we also plan to test the application of GCSplit in eukaryotic datasets and compare it with existing algorithms specialized in either digital normalization or data partitioning.

## Acknowledgments.

This research is supported in part by CNPq under grant numbers 421528/2016–8 and 304711/2015–2. The authors would also like to thank CAPES for granting scholarships. Datasets processed in Sagarana HPC cluster, CPAD–ICB–UFMG.

